# Mitochondrial interactome remodeling in aging mouse skeletal muscle associated with functional decline

**DOI:** 10.1101/2022.06.24.497539

**Authors:** Anna A. Bakhtina, Gavin Pharaoh, Andrew D. Keller, Rudy Stuppard, David J. Marcinek, James E. Bruce

**Author notes:** These authors contributed equally (Anna A. Bakhtina, Gavin Pharaoh).

## Abstract

Genomic, transcriptomic, and proteomic approaches have been employed to gain insight into molecular underpinnings of aging in laboratory animals and in humans. However, protein function in biological systems is under complex regulation and includes factors in addition to abundance levels, such as modifications, localization, conformation, and protein-protein interactions. We have applied new robust quantitative chemical cross-linking technologies to uncover changes muscle mitochondrial interactome contributing to functional decline in aging. Statistically significant age-related changes in protein cross-link levels relating to assembly of electron transport system complexes I and IV, activity of glutamate dehydrogenase, and coenzyme-A binding in fatty acid beta-oxidation and TCA enzymes were observed. These changes showed remarkable correlation with measured CI based respiration differences within the same young-old animal pairs, indicating these cross-link levels offer new molecular insight on commonly observed age-related phenotypic differences. Overall, these system-wide quantitative mitochondrial interactome data provide the first molecular-level insight on ETS complex and substrate utilization enzyme remodeling that occur during age-related mitochondrial dysfunction. Each observed cross-link can serve as a protein conformational or protein-protein interaction probe in future studies making this dataset a unique resource for many additional in-depth molecular studies that are needed to better understand complex molecular changes that occur with aging.

## Main

Aging is a complex process involving several interconnected features that contribute to the progressive decline in function, vulnerability to chronic disease, and ultimately death.^1^ Among the hallmarks of aging is mitochondrial dysfunction which was first proposed as a major component of aging in 1956.^2^ In muscle, aging is accompanied by increasing decline in mass and strength. Decreases in mitochondrial function are thought to be primary mediators of age-related muscle loss.^3^ Many phenotypes of aging have been observed in mitochondria including changes in reactive oxygen species (ROS) production, electron transport system (ETS) efficiency and respiration, ATP production, mitochondrial quality control, mitochondrial biogenesis, and mitophagy.^4^ Mitochondrial function is among the most significant changes accompanying muscle aging on the cellular level.^3,5,6^ Muscle mitochondria have been a primary focus of aging research due to their central role in maintaining metabolic and redox homeostasis, regulating metabolite levels, contribution of mitochondria to muscle function during exercise, and the relative ease of muscle biopsy in humans compared to other tissues. Genomic, transcriptomic, and proteomic approaches have been employed to study muscle aging in laboratory animals and in humans, including deep quantitative proteomic and transcriptomic profiling of several age groups in humans^7,8^ and in mice.^9-11^ However, protein function in biological systems is under complex regulation and includes factors in addition to abundance levels, such as modifications, localization, conformations, and protein-protein interactions. While large-scale studies have also been applied to investigate differential mitochondrial protein modifications with age, including phosphorylation, acetylation, succinylation and others,^12-14^ quantitation of large-scale changes in protein conformations and protein-protein interactions, collectively referred to here as the interactome, has previously not been possible.

Recently developed quantitative crosslinking mass spectrometry technologies (qXL-MS) were applied to elucidate interactome changes in aged murine skeletal muscle mitochondria that contribute to age-related mitochondrial functional decline. Reported here are results from initial investigations of murine muscle mitochondrial interactomes that enable identification of statistically significant changes associated with aging. Newly advanced isobaric quantitative protein interaction reporter (iqPIR) technologies^15^ enabled reproducible detection of age-related mitochondrial interactome changes. Muscle mitochondria from young and old mice were isolated and cross-linked with iqPIR molecules. Young and old mitochondrial samples were paired, processed, and analyzed to quantify age-related mitochondrial interactome changes. Before the cross-linking, mitochondrial protein yield was measured, together with functional measurements such as oxygen consumption rates on Complex I and Complex II substrates, and citrate synthase activity (**Fig. 1**). This allowed the initial correlation to be made linking age-related mitochondrial phenotypic or functional changes with molecular level interactome remodeling. In addition to changes in protein-protein interactions, quantifying site-specific interaction of the iqPIR reporter molecules provides new insights into protein activity by identifying changes in 1) protein structure associated with activity, as in glutamate dehydrogenase, described below and 2) substrate binding to protein active sites. Among these data, significant age-related decreases in cross-link levels within the antenna domain of glutamate dehydrogenase (DHE3) were observed that are correlated with decrease in glutamate and malate driven respiration. Similarly, complex I late-stage assembly and binding of NDUA4 subunit to the rest of complex IV was also impaired and correlated with decrease in complex I respiration. Traditional methods, such as blue native gels (BN-PAGE) are able to distinguish large assemblies, but lack resolution to provide quantitative differences between late stage assemblies.^16^ Moreover, BN-PAGE can only enable visualization of complexes that survive extraction and interactions of ETS complex subunits like NDUA4 appear highly dependent on extraction conditions. Thus, qXL-MS is uniquely suited to study changes in complex assembly and composition and provide new biological insight on ETS dysregulation observed with aging. Finally, as has been previously shown with qXL-MS data,^17^ each identified link can be targeted with PRM methods in other labs to visualize conformational and interactome changes with many other perturbations or interventions. Therefore, in addition to the new biological insight on large-scale age-related protein conformation and interaction changes discussed below, these new quantitative interactome data can serve as a resource for many additional studies to better visualize molecular changes that underpin age-related mitochondrial functional decline.

**Figure 1.**
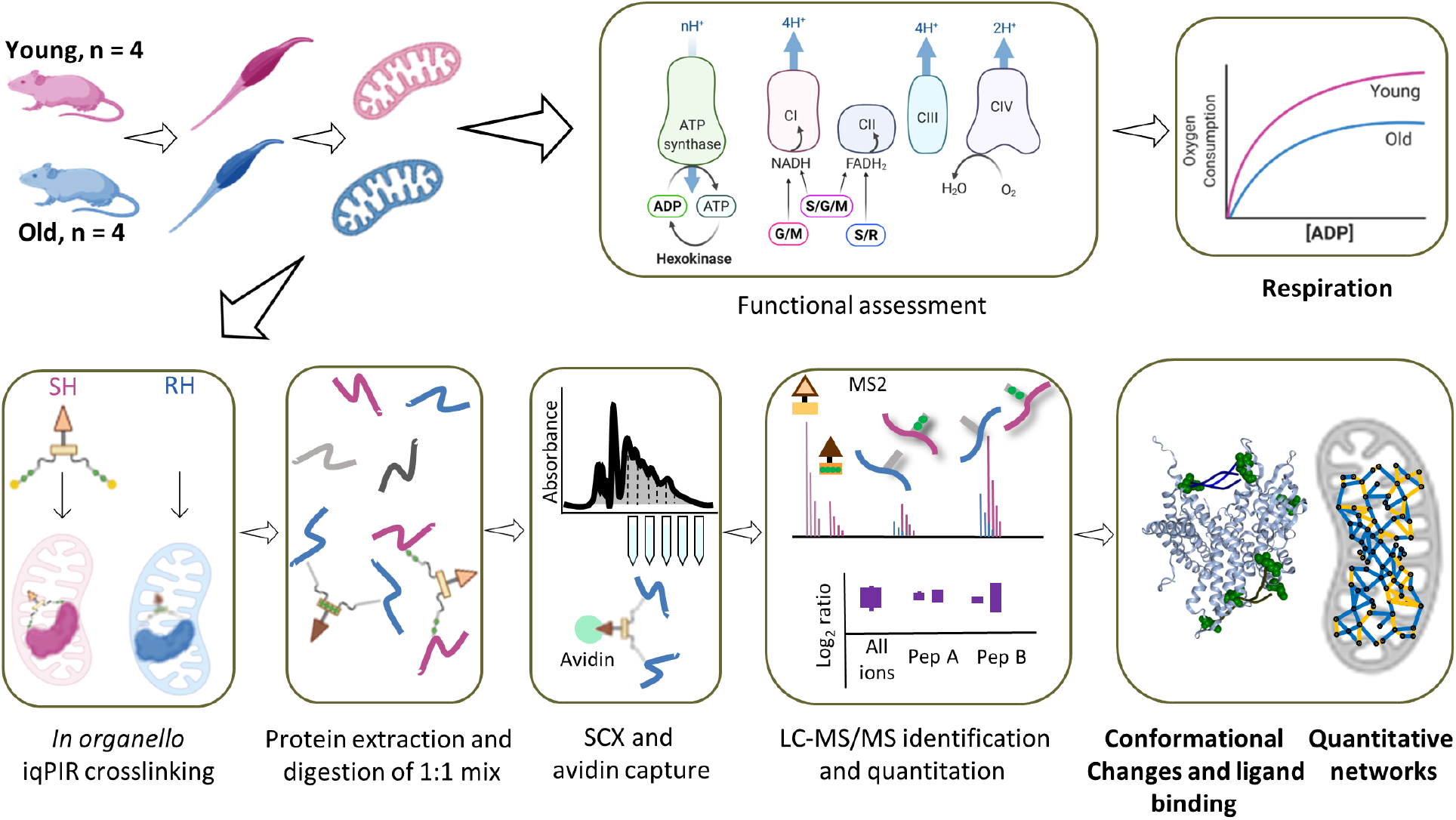
Experimental workflow. Gastrocnemius muscle was excised from either young (6 months) or old (30 months) mice and mitochondria were isolated. Each mitochondrial pellet was resuspended and part of the homogenate was added to an oxygen electrode to measure oxygen consumption for complex I: glutamate and malate (G/M), complex II: succinate and rotenone for inhibiting complex I (S/R), and both complexes: succinate, glutamate, and malate (S/G/M). Mitochondria from the same homogenate were then crosslinked with binary iqPIR reagents: mitochondria from old mice were crosslinked with reporter heavy (RH) and mitochondria from young mice were crosslinked with stump heavy (SH) iqPIR molecules. Mitochondria were then lysed, proteins were reduced, alkylated, and mixed in a 1:1 ratio based on total protein mass for each young old mouse pair and digested with trypsin overnight. Peptide mixtures were then desalted, separated on an SCX column, and enriched for biotinylated peptides with monomeric avidin. Peptides were then separated by LC and MS2 spectra were collected for peptides with charge greater or equal to 4. The data were processed, and abundance of each crosslinked peptide pair was determined using newly developed iqPIR informatics.^15^ The dataset was uploaded to XLinkDB^18^ to view cross-linked peptides, quantitation, protein and complex structures and networks among other dataset features.

## Results

### Generation of mitochondrial interactome of aged muscle

In total, 1864 cross-linked peptide pairs, hereafter referred to as cross-links, were identified from 4 biological replicates of pairwise combinations of 4 old (30 months) and 4 young (6 months) mice at 1% cross-link level FDR (**Supp. Table 1** and http://xlinkdb.gs.washington.edu/xlinkdb/Interactome_of_aged_muscle_mitochondria.php). 533 cross-links are interprotein (formed by lysine residues originating from two distinct proteins) and 1331 are intraprotein (from the same protein). Mapping all identified intralinks on to recently predicted structures^19^ shows that 931 (89%), the overwhelming majority of intralinks, are in agreement with the models: Euclidean distances between alpha carbons of cross-linked lysine residues are less than or equal to 35 angstroms (**Fig. S1a)**. Interlinks are formed between two proximal proteins, so it is expected that these two proteins are localized together within mitochondria. 80% (393) of identified interlinks are between inner membrane associated proteins and only two interlinks are between proteins that are not expected to colocalize (matrix and outer membrane) based on submitochondrial localization information from Mitocarta 3.0 with each pair of interlinked proteins (**Fig. 2a**).^20^

**Figure 2.**
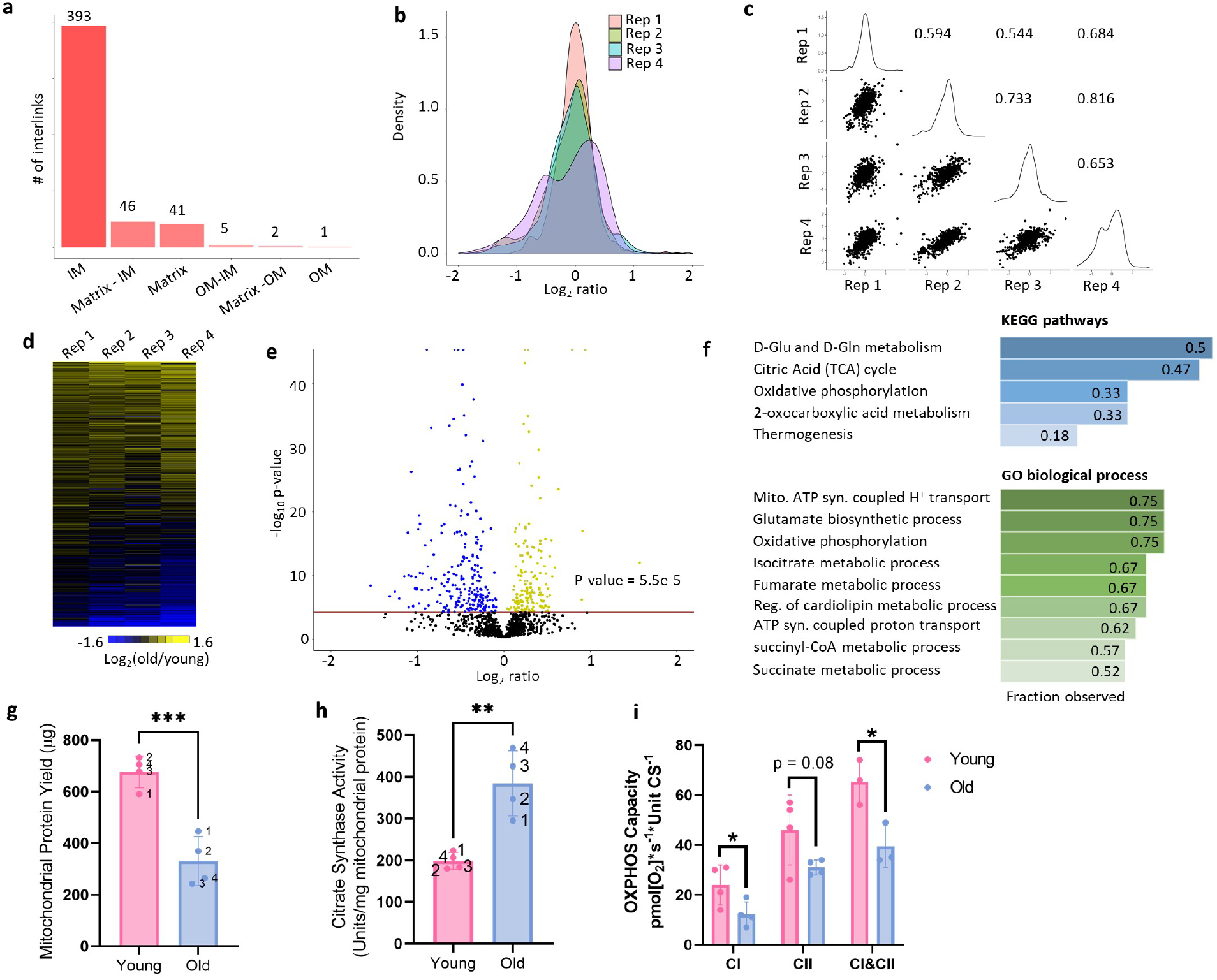
Quantitative cross-linking enables detection of reproducible changes in the interactome of aging mitochondria. **a**. Sub-mitochondrial localization of protein pairs identified in interprotein links. **b**. Distributions of log_2_ ratios for each biological replicate. Pairwise correlation plots (**c**) with Pearson’s R values between 0.54 and 0.81 and heat map (**d**) of log_2_ ratios for cross-links with most reliable quantitation (no missing values and 95% confidence interval < 0.5 for each ratio) show reliable and reproducible quantitation of cross-links. **e**. Confidently quantified cross-links with significant changes (Bonferroni corrected p-value < 0.05 from two-sided t-test). **f**. KEGG and GO Pathways enriched in the cross-links with statistically significant changes. **g**. Yield of mitochondrial protein from both gastrocnemius muscle and (**h**) Citrate Synthase (CS) activity. Each rep of young and old mitochondria is denoted by a number next to the data point. **i**. Maximum oxidative phosphorylation (OXPHOS) capacity of the electron transport chain from isolated mitochondria with complex I (CI) substrates (glutamate and malate), complex II (CII) substrates (succinate and rotenone), or combined CI&CII substrates (glutamate, malate, and succinate) measured as oxygen consumption rate (OCR) at saturating ADP concentrations normalized to units of CS activity. *p < 0.05, **p<0.01, ***p<0.001.

Median normalized log_2_ ratios of confidently quantified crosslinks (no more than 1 missing value across biological replicates and 95% confidence < 0.5) in aged mitochondria compared to young, follow normal distribution, except biological replicate 4 (**Fig. 2b**). To produce log_2_ ratio for a given cross-link, it must be present in both channels in a pair: reporter heavy (RH) cross-link in old sample and stump heavy (SH) cross-link in young. On average, 1156 cross-links are quantified in each replicate (1229, 1099, 1001, 1297 respective), meaning that the majority of the cross-links were formed in both samples, making it unlikely to form by chance. The quantitation derived with iqPIR technologies and informatics showed excellent reproducibility based on observed pairwise Pearson’s R values between 0.5 and 0.76 (**Fig. 2c**). In addition, redundancy in cross-link quantitation exists in some cases because multiple peptide sequences with redundant linkage can be formed during sample processing due to trypsin missed cleavage events during digestion or from methionine oxidation. These cases offer internal quality control on quantitation. Log_2_ ratios for such multiple cross-links generally show excellent agreement (**Fig. S1b**) and further increase confidence in quantitation for a residue pair. Cross-links quantified in every sample show remarkable agreement across the four biological replicates (**Fig. 2d**) and reproducibility and robustness of quantitative values produced by the iqPIR method enabled identification of cross-links that exhibit statistically significant changes (Bonferroni corrected p≤ 0.05) in aging mitochondria (**Fig. 2e**). Analysis of KEGG pathways and GO biological processes show enrichment in proteins involved in glutamate metabolism, TCA cycle, and oxidative phosphorylation (**Fig. 2f** and **Fig. S1g**). As expected, the total mitochondrial protein recovered from each gastrocnemius following isolation was lower in aged samples due to muscle atrophy and decreased input (**Fig. 2g**). However, citrate synthase (CS) activity was significantly increased following mitochondrial enrichment in aged samples (**Fig. 2h**). This finding is consistently reported in the literature, and CS activity is a better metric of mitochondrial content when comparing across ages than mitochondrial protein content^21^ in enriched fractions. Respiration rates in aged mitochondria are significantly decreased when expressed per CS activity (**Fig. 2i, S1f**).

Quantified cross-linked peptide levels are a product of protein, post-translational modifications, conformation and protein interaction level changes. Observed change in a particular cross-link abundance can reflect changes on one or all these levels, unlike traditional proteomics methods that require separate sample preparation, mass spectrometry data acquisition and downstream data processing. We can leverage the entirety of quantified cross-links for each protein or protein pair to make a conclusion regarding which level of regulation is more likely. For example, more than 20 intralinks were quantified from glycerol-3-phosphate dehydrogenase (GPDM) and all are decreased in aged mitochondria (**Fig. S1c**). GPDM is a ubiquinone oxidoreductase which together with its cytosolic counterpart bridges cytosolic energy production and the mitochondrial electron transport chain. High expression of GPDM has been linked to enhanced fatty acid oxidation and resistance to obesity in rats^22^, skeletal muscle regeneration in mice and in cell culture.^23^ GPDM activity can be controlled either by expression levels of the proteins or by allosteric regulation^24^ and consistent decrease in all GDPM cross-links with age most likely indicates lower GDPM levels. Currently no solved mammalian structure of GPDM exists, but the high agreement of cross-links and AlphaFold predicted structure show that cross-linking can be utilized in mechanistic studies for proteins with or without solved structures (**Fig. S1e**). We also observed global decrease of intralinks of MICOS complex proteins and interlinks between its subunits, especially Mic60 and Mic19 (**Fig. S1d**). MICOS complexes establish and regulate cristae morphology and are required for oxidative phosphorylation. The importance of Mic60 and Mic19 and their interaction for MICOS assembly and stabilization has been shown previously and aberrant cristae morphology is a feature of many human pathologies and aging.^25,26^ However, deeper insight regarding age-related conformational and protein interaction changes can be gained from proteins that display specific patterns of change in cross-link levels, allowing for a more detailed analysis.

### Complex I assembly and Complex IV integrity are impaired with age

The present mitochondrial interactome studies resulted in identification of many cross-links originating from electron transport system complexes and supercomplexes (SC) or respirasome interlinks. Respiratory electron transport and complex I biogenesis have been reported as the top pathways affected in aging muscle on a transcriptome level but also the pathways that have the lowest correlation between transcript and protein levels, making the interpretation of its role in aging muscle more complicated.^27^ Decreased NAD^+^/NADH is a hallmark of cell senescence and aging in muscle tissues and is driven in part by CI activity.^28^ CI consists of a membrane embedded part and protruding matrix arm that each assemble independently.^29^ In the matrix arm there are many interlinks as well as intralinks that are either unchanged or slightly increased in aged mitochondria. Conversely, the only matrix arm cross-links with age-related decrease are between NDUS1 and NDUV1 subunits (**Fig. 3 a,b**), yet all intralinks in each subunit are either unchanged or increased (**Fig. 3c**). Comparison of Log_2_ ratios of NDUS1 – NDUV1 cross-links and log_2_ ratios of intralinks from both proteins and their interlinks to other Complex I subunits revealed a statistical difference (p-value = 9.6*10^−6^, Welch two-sample t-test excluding P1 from comparison and 1.35*10^−5^ with all 4 replicates, **Fig. 3d, S2c**). NDUS1 and NDUV1 interlinks with other subunits in the matrix arm and intralinks in all CI subunits except for NDUA8 also do not change or show slight increase with age (**Fig. S2 a, b)**. Two residues on NDUS1 (K84 and K87) were identified cross-linked to the same residue on NDUV1 (K81). One possible explanation for this observation could involve across-link change in post translational modification levels at NDUV1 K81 since it is involved in both links with decreased levels. NDUV1 is a target for desuccinylation by SIRT5^30^ and deacetylation by SIRT3^31^. However, if modification levels at a particular lysine were altered, one would expect that all quantified cross-links involving this residue will change accordingly, indicating a more or less accessible lysine due to the PTM. However, the NDUV1 intra-link (K81-K104) shows no age-related changes, indicating that increased modification of NDUV1 K81 cannot explain the decreased NDUS1-NDUV1 interlinks discussed above. NDUS1 and NDUV1 are among the last subunits to be incorporated in the complex and thus, this interaction is an indicator of fully assembled CI (**Fig. 3e**).^32^ It is also close to flavin mononucleotide (FMN) binding which is a primary site of CI ROS production making these subunits especially vulnerable to ROS damage. The efficient way to replace damaged subunits and keep CI functioning is to replace just the necessary subunits instead assembling the whole complex from scratch. So, the N-module has been shown to have a different turnover rates and mechanisms from Q and P modules. ^33^ Taken together, these data indicate impaired assembly or turnover of N-module intermediate assembly in old mitochondria. Defects in complex I assembly have been shown to lead to increased production of superoxide and premature senescence, and lower abundance of matrix subunits can be a predictor of longevity.^34^ Differential effects of aging on protein abundance and stability of matrix and membrane proteins of Complex I have been previously reported and is expected to lead to impaired assembly.^35^ There have also been recent reports of coordinated assembly of ETC complexes. In particular, complex III was shown to mediate complex I assembly.^36^ CIII deficiency led to stalling of CI assembly, especially incorporation of the N-Module. Interlinks between complex I and complex III as well as several complex III subunits are also decrease in aged mitochondria (**Fig. S2 e, f**).

**Figure 3.**
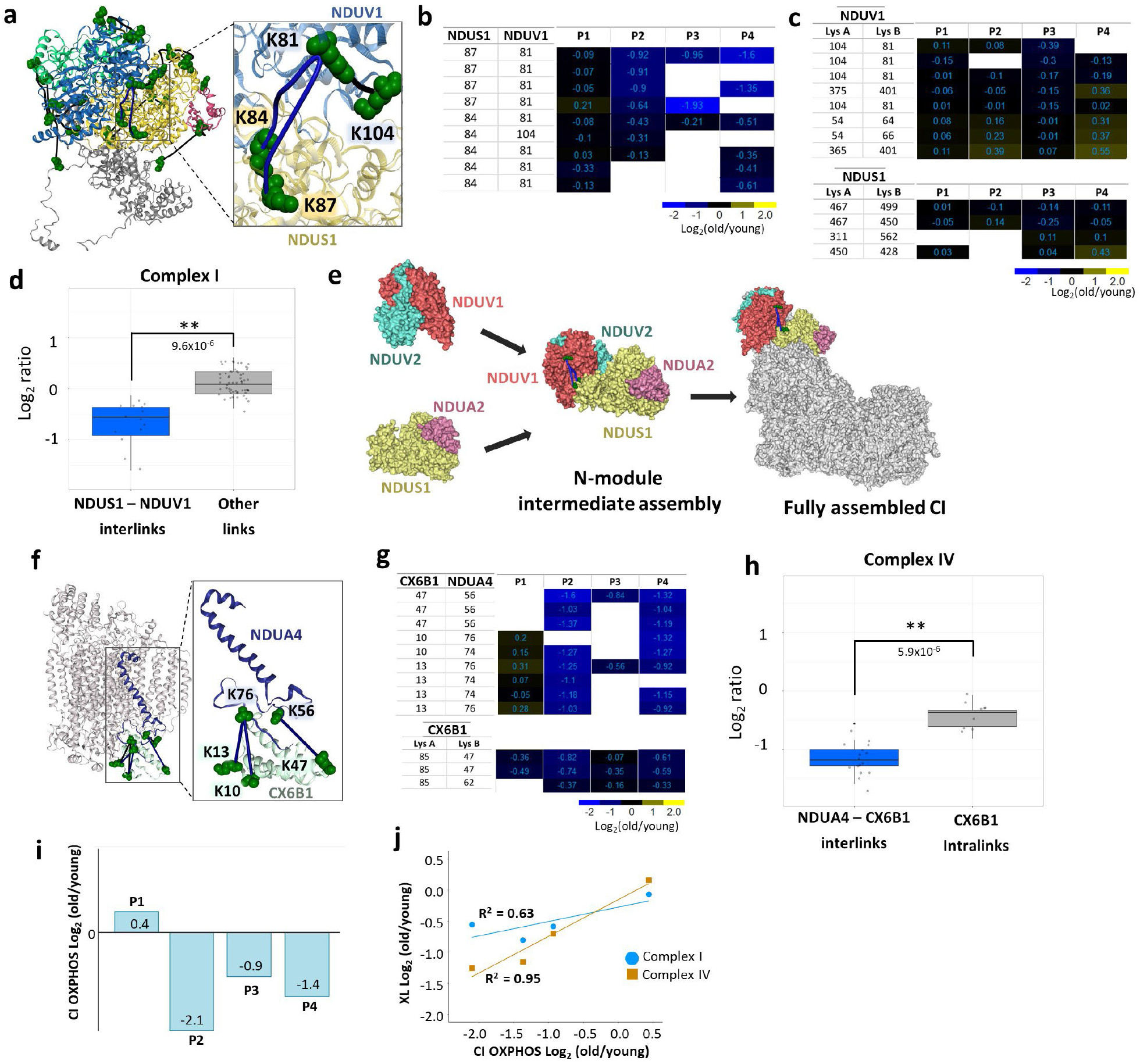
Assembly of Complex I and Complex IV integrity is affected in aging muscle. **a**. Decreased interprotein cross-links between Nduv1 and Ndus1 mapped to Complex I structure (PDB 6G72) are shown in blue and non-changing intralinks are shown in black. **b**. Heatmap of log_2_ ratios of NDUS1-NDUV1 cross-linked peptide pairs for each biological replicate (P1, P2, P3, P4). **c**. Heatmap of log_2_ ratios of intraprotein cross-linked peptides from NDUS1 and NDUV1. **d**. Boxplots of cross-linked peptide pairs in biological replicates P2, P3, P4 with Welch’s t-test p-value < 0.01 show statistically significant difference between NDUS1-NDUV1 interlinks and intralinks and interlinks to other Complex I subunits. **e**. N-module is assembled from subcomplexes NDUV1-NDUV2 and NDUS1-NDUA2 at the end of whole complex I assembly. **f**. NDUA4-CX6B1 interlinks mapped to a CIV structure (PDB 5Z62). **g**. Heatmap of log_2_ ratios of cross-linked peptide pairs between NDUA4 and CX6B1 (top) and CX6B1 intralinks (bottom). **H**. Boxplots of cross-linked peptide pairs in biological replicates P2, P3, P4 with Welch’s t-test p-value < 0.01 show statistically significant difference between NDUA4-CX6B1 interlinks and Cox6b1 intralinks. **i**. Log_2_ fold change of CI linked oxidative phosphorylation in old samples compared to young for each pair. **j**. Correlation plots between average log_2_ ratios of complex I (blue circles) or complex IV (orange squares) cross-links changing with age and log_2_ fold change in Complex I respiration for each pair.

Interlinks between complex IV subunits CX6B1 and NDUA4 were among the cross-links in the dataset that exhibited the largest age-related level decreases(**Fig. 3f, g**). NDUA4 is a small subunit that has been identified to be a subunit of Complex IV rather than complex I as previously thought.^37^ NDUA4 is not required for CIV assembly and CIV is functional without it, but loss of NDUA4 impairs CIV activity.^38^ No NDUA4 intralinks were quantified in this study, but comparing NDUA4-CX6B1 interlinks to CX6B1 intralinks revealed statistically significant decreases of interprotein link levels, p-value = 5.9*10^−6^ excluding P1 from comparison and 0.03 with all 4 replicates (**Fig. 3h and S2d**) indicating reduced interaction between these subunits rather than reduced complex IV levels. Recently, structural characterization of CIV containing NDUA4 subunits has been shown possible by judicious choice of complex extraction/purification conditions^38^ revealing NDUA4 resides at the CIV homodimer interface and precludes CIV homodimer formation.^37^ Moreover, Balsa et al showed that stable knock down of NDUA4 reduced both the activity and stability of CIV that could be rescued by myc-NDUA4 expression.^37^ Therefore, the observed reduction of NDUA4-CIV interaction indicated by reduction in multiple NDUA4-CX6B1 cross-link levels would be expected to decrease CIV stability and activity in mitochondria from old mice. The precise role of NDUA4 and its effect on complex IV is not yet clear and is a subject of ongoing research: a recent report shows replacement of NDUA4 in CIV by NMSE1 (**Fig. S2g)**, another small protein, during inflammation, while NDUA4 is degraded by miRNA.

For both complex I and IV links discussed previously, biological replicate one (P1) deviates from other replicates, showing little or no age-related change in these cross-links (Fig. 3b, g). Overall, CI-linked respiration declined significantly in aged samples (Fig. 2g). Notably however, the aged sample from P1 had no apparent decline in CI-linked respiration compared to young control, which coincides with this pair having reduced changes in CI protein interactions (**Fig. 3i**). Intriguingly, the magnitude of change in the CI assembly cross-links and CX6B1-NDUA4 cross-links in CIV showed strong correlation with the decline in respiration on complex I substrates across all sample pairs: Pearson’s R^2^ 0.63 and 0.95 respectively, while showing no correlation with complex II respiration (**Supp. Table 3** and **Fig. 3j and S2 h**).

### Glutamate dehydrogenase (DHE3) cross-links associated with activation are decreased in aging

Glutamate dehydrogenase (DHE3) is an enzyme responsible for interconversion of glutamate and alpha-ketoglutarate and is encoded by *glud1* gene. A primary DHE3 function *in vivo* is thought to involve catalysis of oxidative deamination of glutamate to produce ammonia and alpha-ketoglutarate.^39^ Alpha-ketoglutarate is a TCA cycle intermediate, but it is also involved in regulation of many cellular processes outside of the TCA cycle, such as epigenetic regulation, protein hydroxylation and ATP synthase regulation (**Fig. 4a**). Connection between DHE3 glycation levels to liver aging has been reported before,^40^ but DHE3 abundance levels have not previously been correlated with aging in muscle. In agreement with that notion, two DHE3 intralink levels were quantified that were unchanged in any of the biological replicates, indicating that the protein abundance levels of DHE3 were not altered with age. Glutamate dehydrogenase exists as a hexamer, comprised of a dimer of trimers, and is a subject of intricate and diverse regulatory mechanisms.^41^ Each trimer forms a protruding structure where helices from all three subunits form an “antenna” with largely unknown function. This antenna is only present in higher organisms and coevolved with the complex regulatory network of DHE3,^42^ suggesting the antenna may serve in a regulatory capacity. Decreased homodimeric links in the DHE3 antenna region were among the largest decreased age-related changes quantified in the present study (**Fig 4b and 3Sa**). These included multiple cross-linked peptide pairs arising from missed cleavage products and were observed in all biological replicates, with more moderated changes in P1 where functional decreases were also moderated (**Fig. 4c, d** and **3Sb**). DHE3 forms an abortive complex upon substrate binding and release of the abortive complex ((DHE3*NAD(P)H*Glu) is facilitated by ADP.^41^ Since ADP serves as an activator of glutamate dehydrogenase, quantitative cross-linking experiments were also performed with mitochondria isolated from HEK293 cells comparing ADP-treated and control untreated mitochondria. These experiments revealed strong increases in DHE3 antenna homodimer links in both biological replicates (**Fig. 4e and 3Se**). An intralink spanning the substrate binding pocket (K171-K352) was also observed with increased level in one of the biological replicates. The combination of ADP-stimulation with old/young interactome data suggests the possibility that DHE3 activity is repressed in aged muscle mitochondria. If so, this may contribute to the observed reduced malate and glutamate stimulated respiration in mitochondria from aged muscle (Fig. 2i). DHE3 is essential for delivery of NADH to complex I during glutamate/malate stimulated respiration and the magnitude of decrease in antenna links correlates with change in Complex I respiration and show no correlation with respiration on complex II (Supp. Table 3 and **Fig. 4f and S3 f**).

**Figure 4.**
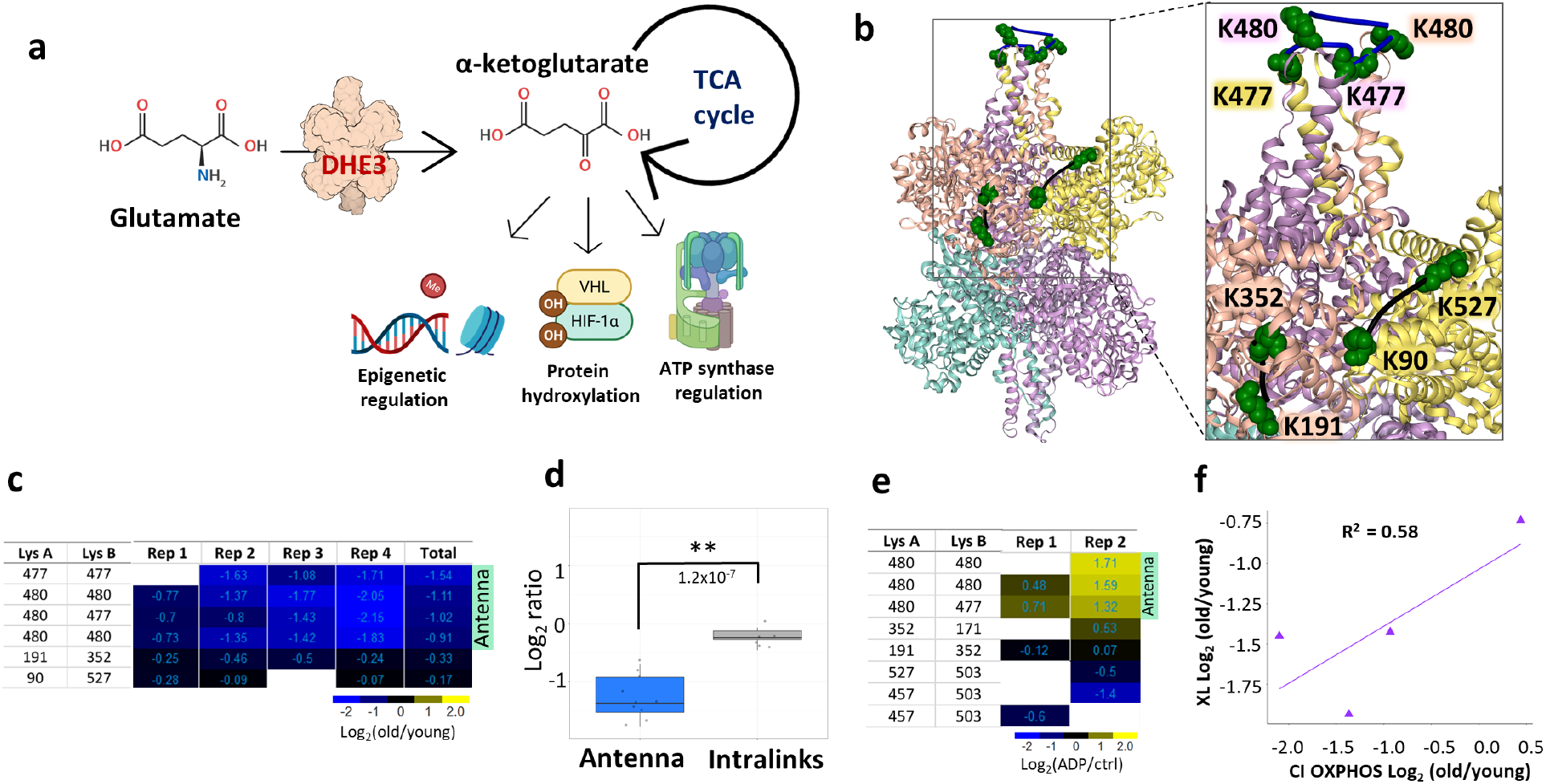
Cross-link levels associated with glutamate dehydrogenase (DHE3) activation are decreased in aged mitochondria. **a**. Glutamate dehydrogenase converts glutamate to α-ketoglutarate, a TCA cycle intermediate that is involved in many cellular processes. **b**. Glutamate dehydrogenase cross-links quantified in aging mouse mitochondria; non-changing intralinks are shown in black and decreasing links in the “antenna” (K477-K477, K477-K480, K480-K480) mapped to one of the trimers in the hexamer (bovine structure 6DHM) are shown in blue. **c**. Log2(old/young) ratio for each cross-linked peptide pair in each biological replicate is summarized in heatmap. **d**. Boxplots of “antenna” ratios and other intralink ratios with Welch two-sided t-test p-value. **e**. Heatmap of cross-linked peptide pairs quantified in 2 biological replicates of ADP treated HEK293 mitochondria. **f**. Correlation between average log_2_ ratio for DHE3 antenna cross-links and CI respiration.

### FAO and TCA cycle enzymes show less accessible substrate binding sites

Impairment of fatty acid metabolism with aging has been shown in multiple organs and models. Aging mouse heart has a decreased free fatty acid flux, TCA cycle flux, and insulin stimulated anaplerosis.^43^ Levels of free fatty acids in blood plasma are decreasing with aging while triglyceride levels are increased.^44^ In addition, muscle contraction leads to a shift in fatty acid oxidation (FAO) and TCA cycle substrate flux and muscle recovery from contraction is impaired with age.^45^ FAO and TCA cycle substrates and intermediates show strikingly different patterns in old mice after the unloading and following recovery compared to young mice. Surprisingly though, transcript levels of the proteins involved do not show significant differences, confounding understanding of the mechanisms underlying the changed FAO and TCA cycle fluxes. Many cross-linked peptides from several FAO and TCA enzymes were quantified in this study, including ACADV, THIL, SUCA and SUCB. ACADV, encoded by *acadvl*, very long-chain acyl-CoA dehydrogenase, catalyzes the first step in beta oxidation (**Fig. 5a**). Although the majority of ACADV Intralinks showed a slight increase in aged mitochondria, indicating possibly slightly elevated protein levels, a subset of four links were significantly decreased in aged muscle mitochondria (**Fig. 5b, h and S4c**). All decreased ACADV links span the binding pocket of fatty acyl-CoA and involve residue K279. The ACADV structure PDB: 2IX5 which contains CoA illustrates that CoA phosphate groups reside within salt bridge formation distance from K279 (Fig. 5A) suggesting that binding of CoA would reduce link formation at this residue. A similar situation was observed with a subset of 8 cross-link levels (out of 18) quantified in the enzyme THIL, encoded by *acat1*, which catalyzes the final FAO step, that showed significant age-related decrease. These link level changes contrast with the remaining 10 THIL cross-links that either slightly increased or showed no change with age. Of the 8 THIL cross-links that decreased with age, 2 span the CoA binding site and K260 which is also within saltbridge distance with CoA phosphate groups as shown (**Fig. 5c, d, i**). Ligand binding can affect cross-link levels within mitochondria as we demonstrated above; both ACADV and THIL age-related decreased links appear statistically enriched in regions involved in ligand binding indicating age-related differences in FAO exist, despite no significant changes in enzyme levels. THIL functions as a tetramer and tetramer formation is linked to higher activity and cancer progression. ^46^ We observed decrease in the homodimeric links indicating decreased levels of active tetramer.

**Figure 5.**
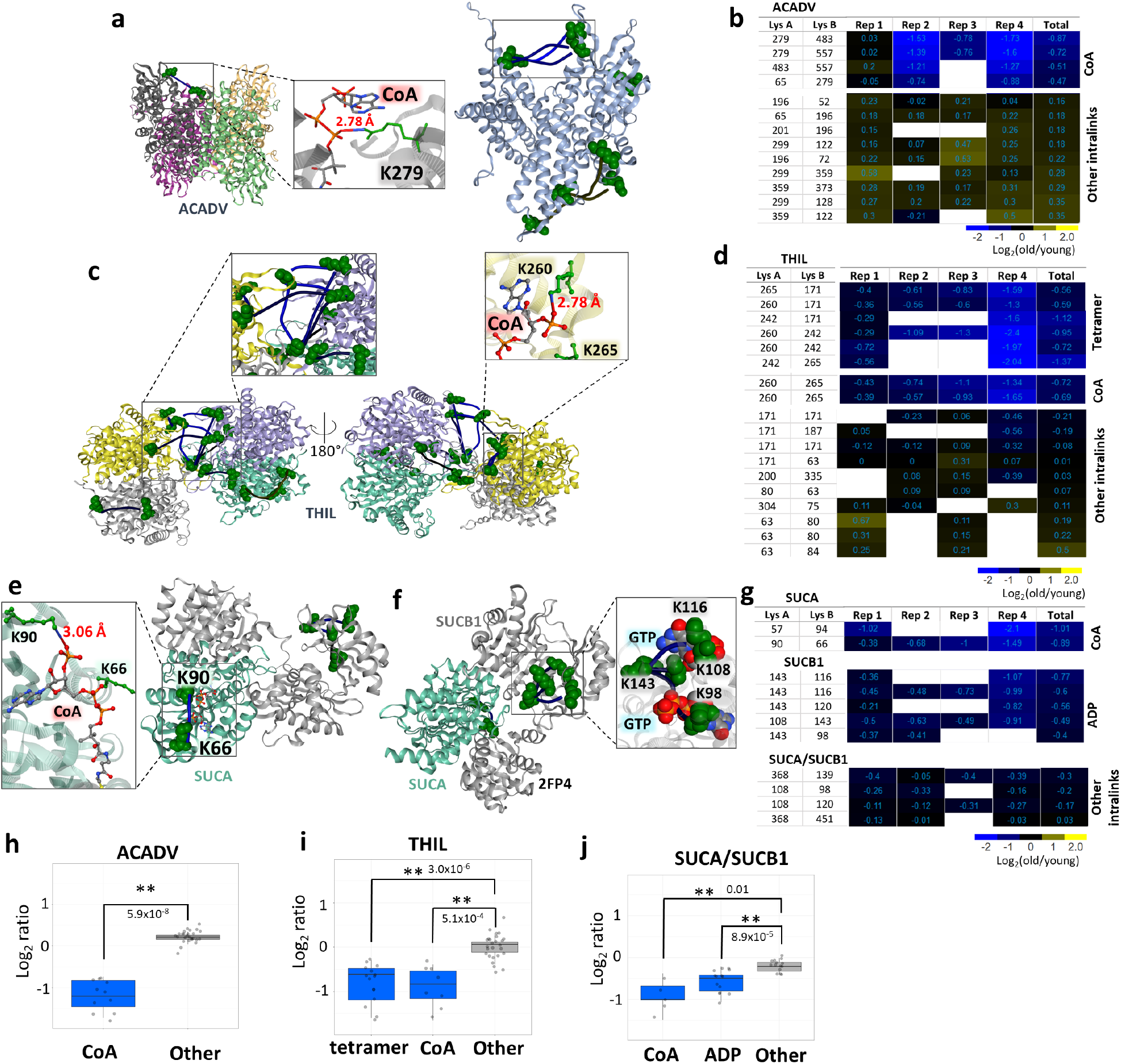
TCA cycle and FA beta oxidation. **a**. ACADV cross-link at the CoA binding site mapped to *A*.*Thaliana* structure, a short chain specific acyl-CoA oxidase in complex with acetoacetyl-CoA (2IX5, left) and all cross-links mapped on a human structure with cross-links at CoA binding site in the box (2UXW, right). CoA from acetoacetyl-CoA is within the distance to form hydrogen bonds with side-chain nitrogen of K279. **b**. Heatmap of log2 ratios of all ACADV cross-linked peptide pairs. **c**. THIL cross-links mapped to a human structure (2IB8) with zoomed in tetrameric links (left). Cross-links at the CoA binding site were also mapped on a human structure crystalized with CoA (2F2S) showing that K260 is within the salt bridge bond formation distance (right zoomed in panel). **d**. Heatmap of log2 ratios of THIL cross-linked peptide pairs. **e**. Succinyl-CoA ligase cross-links mapped on a pig structure of GTP specific succinyl-CoA (4XX0) crystalized with inset view of CoA proximity to SUCA K90. **f**. Succinyl-CoA ligase cross-links spanning ATP/GTP binding site mapped to a pig GTP specific structure (2FP4). **g**. Heatmap of succinyl-CoA ligase cross-links. **h, i, j**. Boxplots comparing decreased cross-link levels at specific sites to log2 ratios of all other intralinks for ACADV, THIL, and SUCA/SUCB1.

The final product of FAO, Acetyl-CoA, then enters the TCA cycle to produce NADH that can then be used by the ETS in oxidative phosphorylation. Significant changes in TCA cycle enzyme cross-link levels in aged mitochondria were also observed, including FUMH (**Fig. S4 a, b**), SUCA and SUCB1. The enzymes SUCA and SUCB1 are subunits of succinate-CoA ligase, which converts succinyl-CoA to succinate that is both a TCA cycle intermediate and an ETS substrate. Both SUCA and SUCB1 exhibited age-related decreased cross-link levels within ligand binding regions, including a cross-link at and nearby SUCA K90 which is the SUCA CoA binding site (**Fig. 5e**). Succinyl-CoA ligase also produces ATP (or GTP in some other tissues) during generation of succinate, and a nucleotide binding pocket exists in SUCB1. A total of 4 intra-links in SUCB1 were observed with age-related decreased levels, all including K143 cross-links that span the nucleotide binding pocket (**Fig. 5f**). Moreover, K108 and K98 in these decreased links exist within a distance compatible with salt bridge formation with GTP phosphate groups as shown in the pig crystal structure (PDB:2FP4). Since other SUCB links appear unchanged with age (**Fig. 5g**), these results indicate that age-related conformation differences within the ligand binding sites and not changes in protein levels mediate the observed changes in succinate-CoA ligase. Indeed, with consideration of the entire group of links, observed ratios of cross-links within both CoA and nucleotide binding regions appear significantly different from those in other succinate-CoA ligase regions (**Fig. 5j**). Although the present age-related changes were measured in young and old murine muscle mitochondria, Wu et al., demonstrated that T cells from rheumatoid arthritis (RA) patients lack sufficient succinyl-CoA ligase activity to maintain balanced TCA cycle metabolic intermediates, implicating acetyl-CoA in controlling pro-inflammatory T cells in autoimmunedisease.^47^ The cross-links identified in the present interactome data offer new opportunities to investigate succinyl-CoA conformational regulation in RA and possibly many other autoimmune diseases in ways not previously possible. Taken together, these results indicate that considerable remodeling of FAO and TCA enzymes occur with aging, yet these appear to not be regulated at the protein level, but rather through conformational differences.

## Discussion

Large-scale capture of quantitative changes in the mitochondrial interactome together with functional measurements provide new molecular insights on age-associated functional decline in bioenergetics and metabolism. While previous transcriptome and proteome studies have provided unparalleled ability to visualize molecular abundance level regulation important in aging, it is clear other regulatory mechanisms beyond protein abundance levels are also involved. The approach presented here combines quantitation of protein, conformation, modification, ligand binding, and protein interaction levels to provide new biological insight on age-related molecular changes. Recently, the importance of studying protein interaction and their role in aging has been brought to the community attention.^48^ While this initial study is non-comprehensive, these efforts have yielded the largest quantitative interactome dataset to define age-related mitochondrial differences thus far and include 1521 quantified cross-links.

To date, changes in glutamate dehydrogenase mRNA or protein levels with aging have not been reported and the studies presented here are consistent with that finding. However, the present quantitative cross-linking data generate new insights on DHE3 interactions in mitochondria and age-related conformational differences that may functionally contribute to age-associated changes connecting TCA cycle, ETS and alpha-ketoglutarate (aKG) effects on lifespan. Multiple reports demonstrate aKG involvement in lifespan extension in mice^49^, flies^50^, yeast^51^ and worms.^52^ Increase in glutamate dehydrogenase activity has also been shown to accompany caloric restriction and subsequent increased lifespan.^53^ Moreover, diet-based lifespan extension in flies appears to be dependent on DHE3 expression.^54^ Alpha-ketoglutarate also promotes myofibroblast differentiation through epigenetic regulation by driving histone demethylation and the role of anaplerotic supply from GLN and GLU has been highlighted.^55^ The decreases in DHE3 antenna cross-link levels presented here are not resultant from protein level changes and indicate PTM and or conformational differences exist in aged muscle mitochondria. Age-related increase in PTM levels is possible since both cross-linked residues, K477 and K480 are targets of the sirtuins SIRT3 and SIRT5, with acetylation levels of these residues increasing more than 8 fold upon SIRT3 knock out.^30,31,56^ However, cross-linked sites in the DHE antenna show both decrease levels in aged vs young mitochondria where glutamate respiration is repressed, and increased levels in ADP stimulated mitochondria where glutamate dehydrogenase activity is increased. In addition to the decrease in maximum glutamate stimulated complex I respiration, we also observed decreased maximum glutamate stimulated respiration and lower sensitivity of respiration to glutamate in aged male CB6F1 mice compared to young. (**Fig. S3 c, d**). This suggests that disrupted glutamate metabolism and the role of glutamate dehydrogenase is not strain or sex specific. Therefore, these antenna cross-link levels can serve as probes of glutamate dehydrogenase activity in many future studies, including those to help unravel Sirtuin-, diet-, or exercise-mediated lifespan or healthspan extension. For instance, quantitation of DHE3 antenna links prior to age-induced changes in mitochondrial function, with caloric restriction or other interventions can help elucidate pathways that mediate age-related reduction in glutamate respiration.

Quantitative cross-linking revealed changes in protein interactions and conformations affecting many facets of metabolism in aging muscle. Increased ROS production and alterations among ETS complexes in mitochondria are among the primary aspects under study to better understand age-related mitochondrial functional decline. ETS complexes, especially complex I, require coordinated and controlled assembly to achieve functional maturity.^57^ Therefore, disruption of the assembly uncovered in the present study should be investigated further to elucidate its role in ETC pathologies in aging. Altered activity and ligand binding in FAO and TCA cycle enzymes can bring to the forefront the contributions of TCA intermediates and fatty acid metabolism to aging phenotypes, connecting phenotypes and molecular remodeling. ^58-60^ Excitingly, we have already observed decreased sensitivity to fatty acids in the aged CB6F1 mice similarly to decreased sensitivity to glutamate (**Fig. S4 d, e**).

Taken together, these data enable a system-wide view of the changing mitochondrial interactome landscape linking changes in glutamate dehydrogenase activity together with amino acid metabolism, TCA cycle, and energy production by oxidative phosphorylation (**Fig. 6a**). All the age-related changes highlighted in this figure involve changes in protein conformations and interactions that are not readily attainable through conventional protein abundance level quantitation. Strikingly, many age-related interactome changes appear well-correlated to the severity of aging mitochondrial phenotype, as shown with pairwise CI oxygen consumption ratio compared with the magnitude of changes in protein conformations, interactions and ligand binding (**Fig. 6b)**.

**Figure 6.**
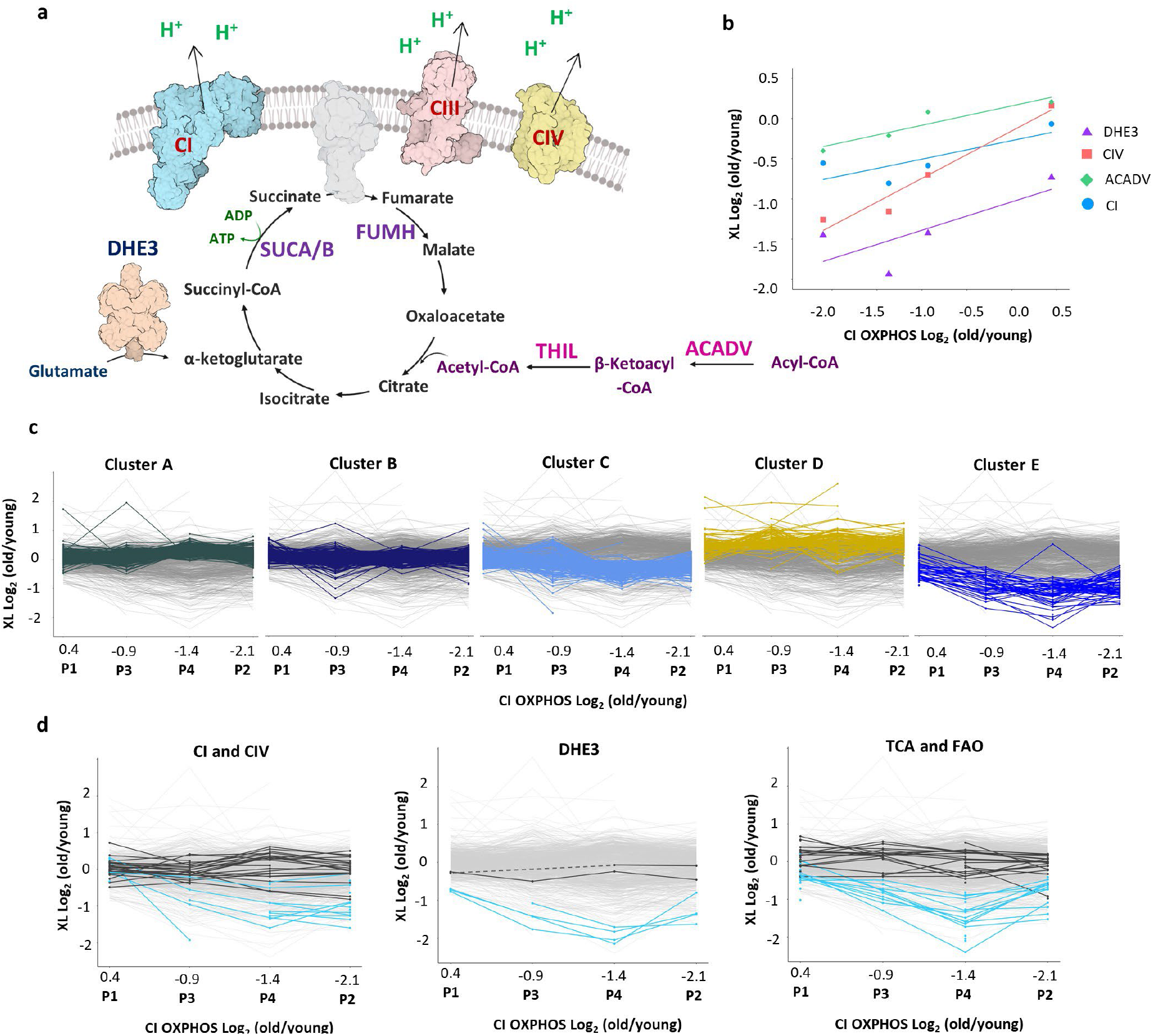
Interactome remodeling associated with changes in muscle metabolism with aging. **a**. Integrated pathways with age-associated changes in protein-protein interactions, protein-ligand interactions, or conformational changes highlighted in this study. **b**. Correlation between average of log_2_ ratios in each biological replicate of cross-links changed with age in DHE3 (purple triangles, R^2^=0.58), Complex IV (red squares, R^2^=0.95), ACADV (green rhombi, R^2^=0.87), Complex I (blue circles, R^2^=0.63) and Complex I driven respiration. **c**. K-means clustering with 5 clusters. **d**. Crosslinks with age-related changing levelsthat are discussed in this study (blue) cluster together (cluster E); non-changing crosslinks from the same proteins are in dark grey

In the present manuscript, detailed discussion of only a small subset of cross-links was possible. K-means cluster analysis of all cross-links quantified in at least 2 biological replicates (95% conf. =<1) revealed that cross-links correlating with functional measurements cluster together and non-changing cross-links from these proteins are in a separate cluster together (**Fig. 6c, d**). Many other cross-links in these proteins display similar patterns to those discussed above with functional measurements, including proteins from the same pathways, such as ACSL1, CMC1, CPT1B, ATPB, and ATP5F1. The entire interactome dataset with quantitation, structures with mapped cross-links, and k-means clustering assignments is available to view online in XLinkDB (http://xlinkdb.gs.washington.edu/xlinkdb/Interactome_of_aged_muscle_mitochondria.php). These data provide a unique, detailed, and quantitative view of mitochondrial aging in muscle that can be used to guide future studies unraveling molecular underpinnings of metabolism changes with age.

## Methods

### Animal Husbandry

This study was reviewed and approved by the University of Washington Institutional Animal Care and Use Committee. Female 6-month and 30-month-old C57BL/6J mice were received from the Jackson Laboratory. All mice were maintained at 21 °C on a 14/10 light/dark cycle and given standard mouse chow and water ad libitum with no deviation prior to or after experimental procedures. C57BL/6J animals were killed by cervical dislocation with no anesthetic. CB6F1 mice were euthanized with beuthanasia.

### Mitochondrial Isolation

The gastrocnemius muscle was dissected, and mitochondrial isolation was performed by differential centrifugation. The whole muscle was homogenized using a high-speed drill on ice in a glass Dounce homogenizer in Mitochondria Isolation Buffer (210 mM Sucrose, 2 mM EGTA, 40 mM NaCl, 30 mM HEPES, pH 7.4). The homogenate was centrifuged at 900 x g at 4 °C for 10 minutes. The supernatant was collected and centrifuged at 10,000 x g at 4 °C for 10 minutes. The supernatant was removed, and the mitochondrial pellet was resuspended in ice-cold Respiration Buffer (RB) without taurine or bovine serum albumin (BSA) (1.5 mM EGTA, 3 mM MgCl_2_-6H_2_O, 10 mM KH_2_PO_4_, 20 mM HEPES, 110 mM Sucrose, 100 mM Mannitol, 60 mM K-MES, pH 7.1). The respiration buffer for mitochondrial resuspension did not include taurine, because it is an aminoethane sulfonic acid which contains a primary amine that could react with the cross-linker. F1 mice resuspension media had taurine. Isolated mitochondria protein concentration was determined using standard Bradford Assay procedures.

### Mitochondrial Respiration

CI, CII, and CI&CII-linked mitochondrial respiration were assayed in mitochondria isolated from young (6-mo-old) and old (30-mo-old) female C57Bl6/J mouse gastrocnemius using an Oxygraph 2K dual respirometer/fluorometer (Oroboros Instruments, Innsbruck, Austria). RB with taurine and BSA was used for respiration measurements (1.5 mM EGTA, 3 mM MgCl_2_-6H_2_O, 10 mM KH_2_PO_4_, 20 mM HEPES, 110 mM Sucrose, 100 mM Mannitol, 60 mM K-MES, 20 mM taurine, 1 g/L BSA, pH 7.1). Hexokinase clamp (1 U/ml hexokinase, 2.5 mM D-glucose) was used to maintain equilibrium of ATP/ADP at submaximal ADP concentrations.^61^ Respirometry and fluorometry reagent stocks were prepared according to Oroboros instructions (bioblast.at). Respiration was measured at 37°C with stirring during substrate and inhibitor titrations.

To measure CI, CII, and CI&CII-linked respiration, first, 10 μM cytochrome c was added to each chamber to allow measurement of respiration in isolated mitochondria without limiting by membrane damage occurring during isolation. Approximately 35 μg mitochondrial homogenate (∼8-11 μL) was added to each 2 mL chamber. Complex I (CI), Complex I (CII), and CI&CII-linked respiration were measured in parallel for each sample by adding complex-specific substrates and inhibitors then titrating in ADP. CI-linked respiration was measured by adding 10 mM glutamate and 0.5 mM malate. CII-linked respiration was measured by adding 10 mM succinate and 0.5 μM rotenone. CI&CII-linked respiration was measured by adding 10 mM succinate, 10 mM glutamate, and 0.5 mM malate. The OXPHOS capacities for each substrate condition were determined as the maximum oxygen consumption rate (OCR) measured during a titration of ADP from 5-6000 μM ADP. The background oxygen consumption with de-energized mitochondria was subtracted from all measured functional parameters before reporting final values.

Response to glutamate and fatty acid titration was measured in mitochondria isolated from 8 young (5-7-mo-old) and 6 old (33-37-mo-old) male CB6F1 mouse gastrocnemius using the Oxygraph 2K dual respirometer/fluorometer. RB with taurine and BSA was used for respiration measurements without hexokinase clamp because saturating ADP concentrations were added to the chambers in a single bolus during the experiment. To measure glutamate sensitivity, a sequential titration of 50 μg mitochondrial protein, 2.5 mM ADP, 10 μM cytochrome c, and sequential additions of 1 mM glutamate up to 10 mM glutamate final concentration were performed. To measure fatty acid utilization, a sequential titration of 50 μg mitochondrial protein, 2 mM malate, 2.5 mM ADP, 10 μM cytochrome c, and sequential additions of 10 μM palmitoyl-carnitine up to 100 μM palmitoyl-carnitine final concentration were performed.

The mitochondrial respiration results were analyzed using Microsoft Office Excel and GraphPad Prism 9.9 for Mac OS X (GraphPad Software, La Jolla, CA). For all comparisons, *P* < 0.05 was considered statistically significant. Comparisons between two groups were analyzed using unpaired two-tailed student’s t-test. Comparisons during ADP titrations were analyzed using repeated measures Two-way ANOVA with Sidak’s multiple comparisons. Comparisons during glutamate and palmitoyl-carnitine titrations in CB6F1 mice were analyzed using mixed-effects analysis with Sidak’s multiple comparisons. Plots depict mean ± standard deviation

### Citrate Synthase Activity Assay

Citrate Synthase (CS) activity is reportedly a more accurate marker of mitochondrial mass than total protein content when performing comparisons across age^21^. CS activity assay was performed on mitochondrial isolations and used to normalize mitochondrial respiration. CS Activity was measured by spectrometric quantitation (412 nm) of 5,5’dithiobis-2-nitrobenzoic acid conversion to 2-nitro-5-thiobenzoic acid in the presence of Coenzyme A thiol generated during citrate production (CS0720, Sigma) as previously described.^62^

### Cross-linking of isolated muscle mitochondria

Isolated mitochondria from murine gastrocnemius muscles of 8 mice (4 young and 4 old) were resuspended in cross-linking buffer (170 mM Na_2_HPO_4_, pH 8.0) and either reporter heavy (RH) or stump heavy (SH) iqPIR reagent was added;^15^ final reaction volume was 100 uL and cross-linker concentration was 10 mM. Cross-linking reaction was allowed to proceed for 30 min at room temperature with shaking. Cross-linking buffer was then removed by centrifugation and mitochondrial pellets were lysed in 8M urea. Proteins were reduced with TCEP (30 min RT with shaking) and alkylated with IAA (30 min RT with shaking). Protein concentration of each mitochondrial sample was measured with a Bradford assay using Cytation plate reader. Samples were mixed pairwise (one old and one young, **Supp. Table 4**) using equal amount of protein from each sample making 4 biological replicates total. Protein mixtures were digested with trypsin overnight (1:100 trypsin concentration at 37 C with shaking). Peptides were then acidified with TFA and cleaned using seppak c18 columns (Waters). Peptides were separated using SCX chromatography (Luna column, Agilent HPLC) into 14 fractions and fractions were pooled together as following: fractions 1 to 5, fractions 6 and 7, fraction 8, fraction 9, fraction10, fractions 11 to 14. Pooled fractions were dried in a SpeedVac and resuspended in ammonium bicarbonate buffer; pH was adjusted to 8.0 with NaOH. Biotinylated cross-linked peptides were captured with monomeric avidin (ThermoFisher Scientific 20228) for 30 min at RT with shaking. The beads were washed with ammonium bicarbonate and peptides were eluted with 0.1% formic acid in 70% ACN, dried down by vacuum centrifugation and resuspended in 20 uL of 0.1% formic acid.

### Mitochondrial isolation from HEK293 cells and treatment with ADP

HEK293 cells were grown in DMEM media supplemented with 3.5 mg/L glucose, 10% fetal bovine serum, 1% penicillin and streptomycin to confluency. The plates were washed with PBS, cells detached using EDTA 20 mM, centrifuged and washed twice in MgCl_2_. Cells were then resuspended in ice-cold mitochondrial isolation buffer (70 mM sucrose, 220 mM D-mannitol, 5 mM MOPS, 1.6 mM carnitine, 1 mM EDTA at pH 7.4) and homogenized in a glass homogenizer. The homogenate was centrifuged at 600 g for 5 min at 4 C. The supernatant was transferred to a 15 mL tube and centrifuged at 8000 g for 10 min at 4 C. The supernatant was then removed, and mitochondrial pellet was resuspended in 5 mL of mitochondrial isolation buffer and centrifuged at 8000 g for 10 min. The mitochondrial pellet was then resuspended in 200 uL of mitochondrial isolation buffer and split into two. ADP was added to one vial to a final concentration of 1.5 mM. Both samples were incubated at RT for 10 min with shaking. Supernatant was then removed by centrifugation and pellets were resuspended in cross-linking buffer. RH iqPIR cross-linker and ADP was added to ADP treated sample to final concentrations of 10 and 1.5 mM respectively. SH iqPIR cross-linker was added to control sample to a final concentration of 10 mM. The cross-linking reaction was allowed to proceed for 30 min at RT with shaking. The supernatant was then removed by centrifugation and mitochondrial pellets were lysed, reduced, alkylated, combined, digested and processed for mass spectrometric analysis the same way as murine muscle mitochondria.

### Mass Spectrometry and data analysis

Four uL of each pooled fraction was loaded on a 60 cm C8 heated column and separated on 2 hour gradient on nanoAcquity HPLC system (Waters) and analyzed with QExactive Plus mass spectrometer (ThermoFisher Scientific). MS1 scans were analyzed at 70K resolution with AGC target 1e6, and maximum ion time 100 ms. Top 5 peaks with charge 4 or greater were selected for HCD fragmentation with NCE 30 and MS2 spectra were collected at 70K resolution, 5e4 AGC target, and 300 ms maximum ion time.

Raw files were converted to mzXML, and spectra containing cross-linked peptides were determined with Mango software.^63^ These spectra were then searched against mouse Mitocarta 2.0 database using Comet^64^ search engine and cross-linked peptides were validated with XLinkProphet.^65^ Identified cross-links were quantified using iqPIR algorithm and results were uploaded to XLinkDB database.^18^ Normalized log_2_ ratios and associated p-values based on the Student’s t-test on each quantified ion for every cross-link (t = sqrt(df*mean/std) and p-values calculated with the pt function of R: pt(-abs(t), df) where t is the t-statistic and df the degrees of freedom) were downloaded from XLinKDB and correlation plots between biological replicates, density plots for each replicate, volcano plot indicating significantly changed cross-links, box-plots and t-test comparisons were generated in R using tidyverse package and R markdown is provided.^66^ In all boxplots horizontal line represents median, the lower and upper hinges correspond to the first and third quartiles (the 25th and 75th percentiles) and the whiskers extend to the value no further than 1.5 IQR. Pathway enrichment analysis and network of differentially expressed cross-links were generated using STRING web-application.^67^ Heatmap of all common cross-links was generated for cross-links quantified with 95% confidence (interval within which one can be sure with 95% confidence that the actual mean value resides, calculated as 1.96 * std / sqrt(num_reps) assuming normal distribution) less than 0.5 in all 4 biological replicates using NG-CHM builder web application.^68^ Heat maps for cross-links in specific proteins or protein complexes were generated in XLinkDB. Cross-links were mapped on available structures with either Euclidean distances or SASD distances calculated by Jwalk.^69^

## Abbreviations

ETS: Electron Transport System
TCA: Tricarboxylic acid
FAO: Fatty Acid Oxidation
LC-MS: Liquid Chromatography – Mass Spectrometry
ROS: Reactive Oxygen Species
ATP: Adenosine Triphosphate
qXL-MS: quantitative crosslinking mass spectrometry
iqPIR: isobaric quantitative protein interaction reporter
SCX: Strong Cation Exchange
SH: Stump Heavy
RH: Reporter Heavy
GO: Gene Ontology
KEGG: Kyoto Encyclopedia of Genes and Genomes
CS: Citrate Synthase

## Data and code availability

Mass spectrometry data have been deposited to ProteomeXchange Consortium via PRIDE repository with identifiers PXD031643 and PXD031644. R markdown used for statistical analysis and figure generation is available at https://github.com/brucelab/aging_mito_interactome_data_analysis.

## Acknowledgments

Figures were created with BioRender.com. Authors are grateful to members of the Bruce lab for helpful discussion. This work was funded by NIH grants P01-AG001751, R56-AG070096, T32 AG066574, R35GM136255, and R01HL144778.

## Extended Data

**Supplemental Table 3.** Log_2_ fold change calculated for each paired biological replicate in mitochondrial yield, citrate synthase (CS) activity, ad oxygen consumption on either CI substrates or CII substrates.

| Biorep | Mito yield | CS activity | CI oxphos | CII oxphos |
| --- | --- | --- | --- | --- |
| P1 | -0.712 | 0.712 | 0.441 | -0.667 |
| P2 | -0.878 | 0.918 | -2.1 | -0.788 |
| P3 | -1.282 | 0.932 | -0.933 | 0.107 |
| P4 | -1.417 | 1.186 | -1.369 | -0.697 |

**Supplemental Table 4.** Pairwise combinations of old and young mitochondria to create 4 biological replicates (mice numbered the same way as in functional data). Pairs were assigned based on amount of available protein to maximize input material.

| Biorep | Old mouse # | Young mouse # |
| --- | --- | --- |
| P1 | 1 | 2 |
| P2 | 2 | 3 |
| P3 | 3 | 1 |
| P4 | 4 | 4 |

**Extended data figure S1.**
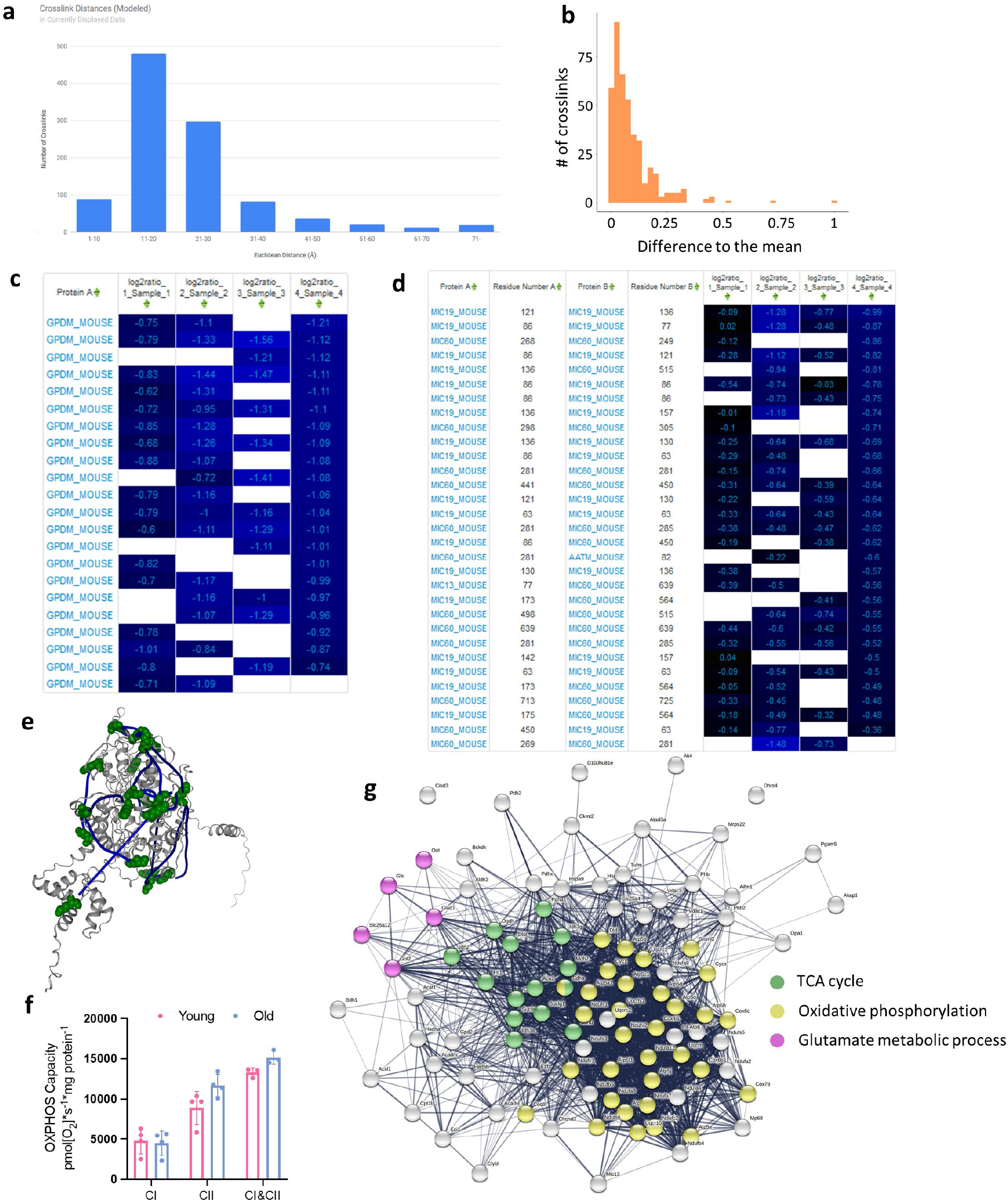
**a** Histogram of calculated Euclidean distances for all intraprotein cross-links mapped to AlphaFold predicted structures. **b**. Histogram of differences between a mean log2 ratios for cross-linked residue pairs based on multiple cross-linked peptides (cross-links that connect the same lysines, but can be identified in differently cleaved or modified peptides) and each cross-linked peptide pair **c**. Heatmap of log2 ratios of GPDM Intralinks in 4 biological replicates. **d**. Heatmap of log2 ratios of MICOS complex subunits intra- and interprotein links. **e**. GPDM Intralinks mapped to an AlphaFold predicted structure. **f**. Oxphos capacity normalized by protein amount. **g**. STRING network of proteins with significantly changed cross-links.

**Extended data figure S2.**
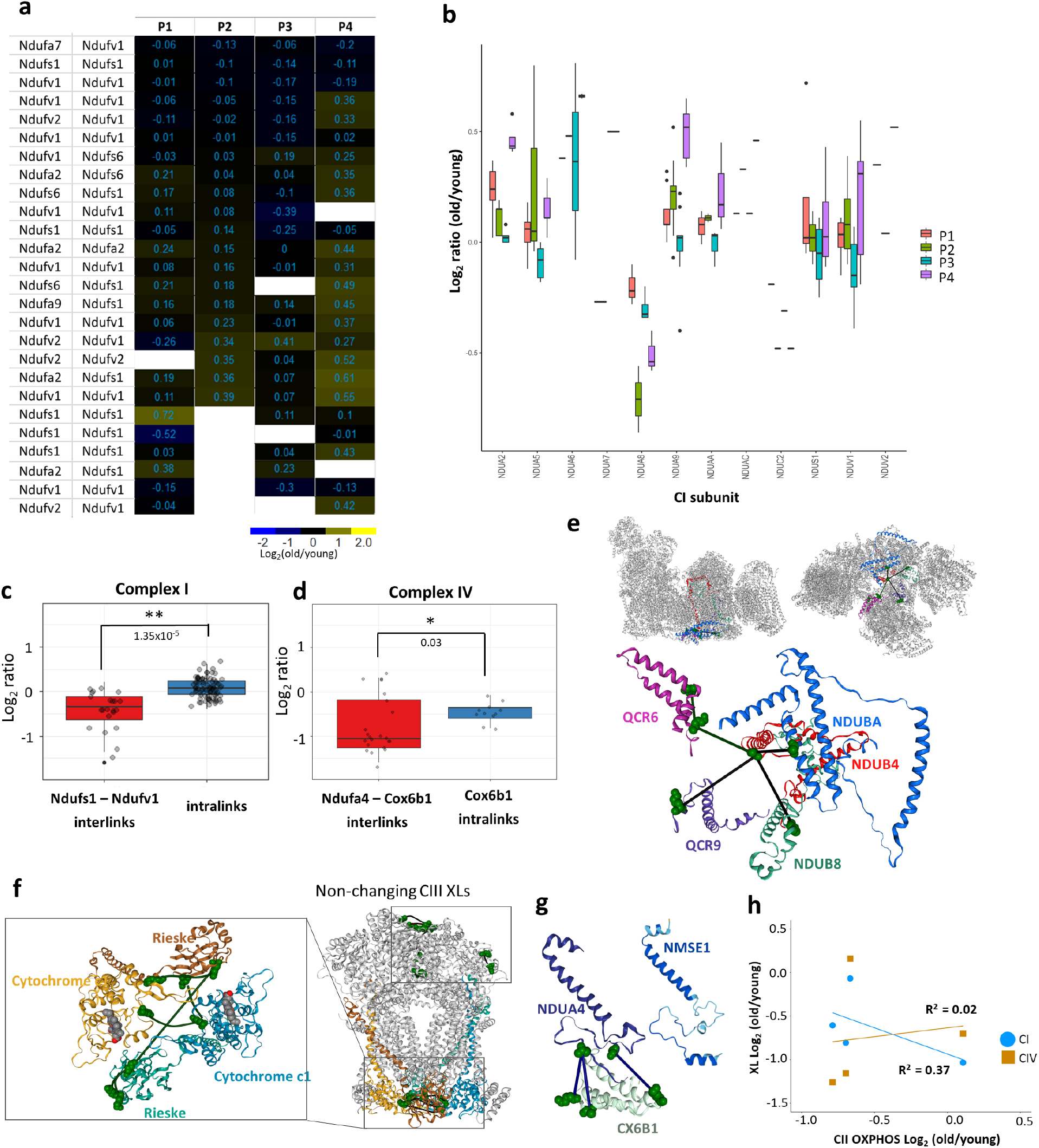
**a**. Heatmap of log2 ratios of all NDUS1 and NDUV1 crosslinks cross-links and interprotein crosslinks to other CI subunits. **b**. Boxplots of all intralinks in Complex I subunits by a biological replicate. **c**,**d**. Boxplots of crosslinks downregulated in aging and non-changing Intralinks based on all 4 biological replicates for CI and CIV respectively. P-values are from Welch t-test and significance (** for 0.01 and * for 0.05). **e**. Structure of supercomplex with cross-linked CI and CIII subunits highlighted (top) and specific CI-CIII crosslinks mapped to the subunits (bottom); decreased crosslinked are in green. **f**. Complex III crosslinks mapped to a bovine structure. Subunits with decreased intralinks highlighted and zoomed in (right). **g**. AlphaFold predicted structure of human NMSE1, that is reported to replace NDUA4 subunit in complex IV during inflammation, and cryo-EM structure of CX6B1 and NDUA4 in CIV. **h**. Correlation plots of CI and CIV cross-links changing in aging mitochondria and Complex II OXPHOS.

**Extended data figure S3.**
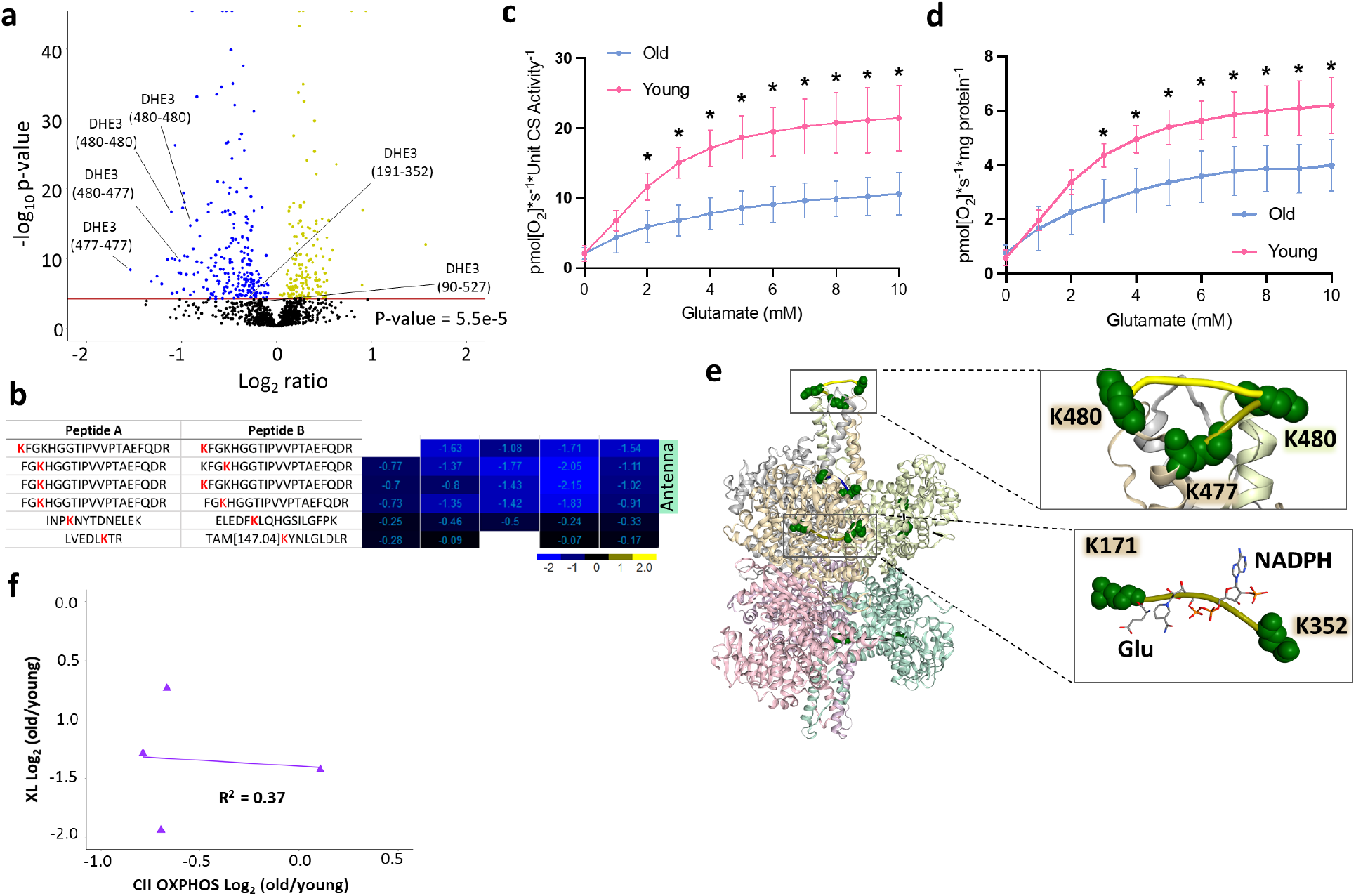
**a**. Decreased and non-changing cross-link levels in glutamate dehydrogenase highlighted on the volcano plot with Bonferroni corrected p-value = 0.05. **b**. Heatmap of all DHE3 cross-linked peptide pairs with each individual peptide sequence shown. Cross-linked lysine residues are in red. **c, d**. Glutamate sensitivity assay as measured by oxygen consumption determined with glutamate titration and normalized by citrate synthase activity or mitochondrial protein amount. **e**. Antenna cross-links increased with activation by ADP treatment of isolated HEK293 mitochondria mapped to 6DHM structure. Cross-link spanning active site (K171-K352) shows slight upregulation indicating possible destabilization of abortive complex by ADP. **f**. Correlation plots of DHE3 antenna cross-links changing in aging mitochondria and Complex II OXPHOS.

**Extended data figure S4.**
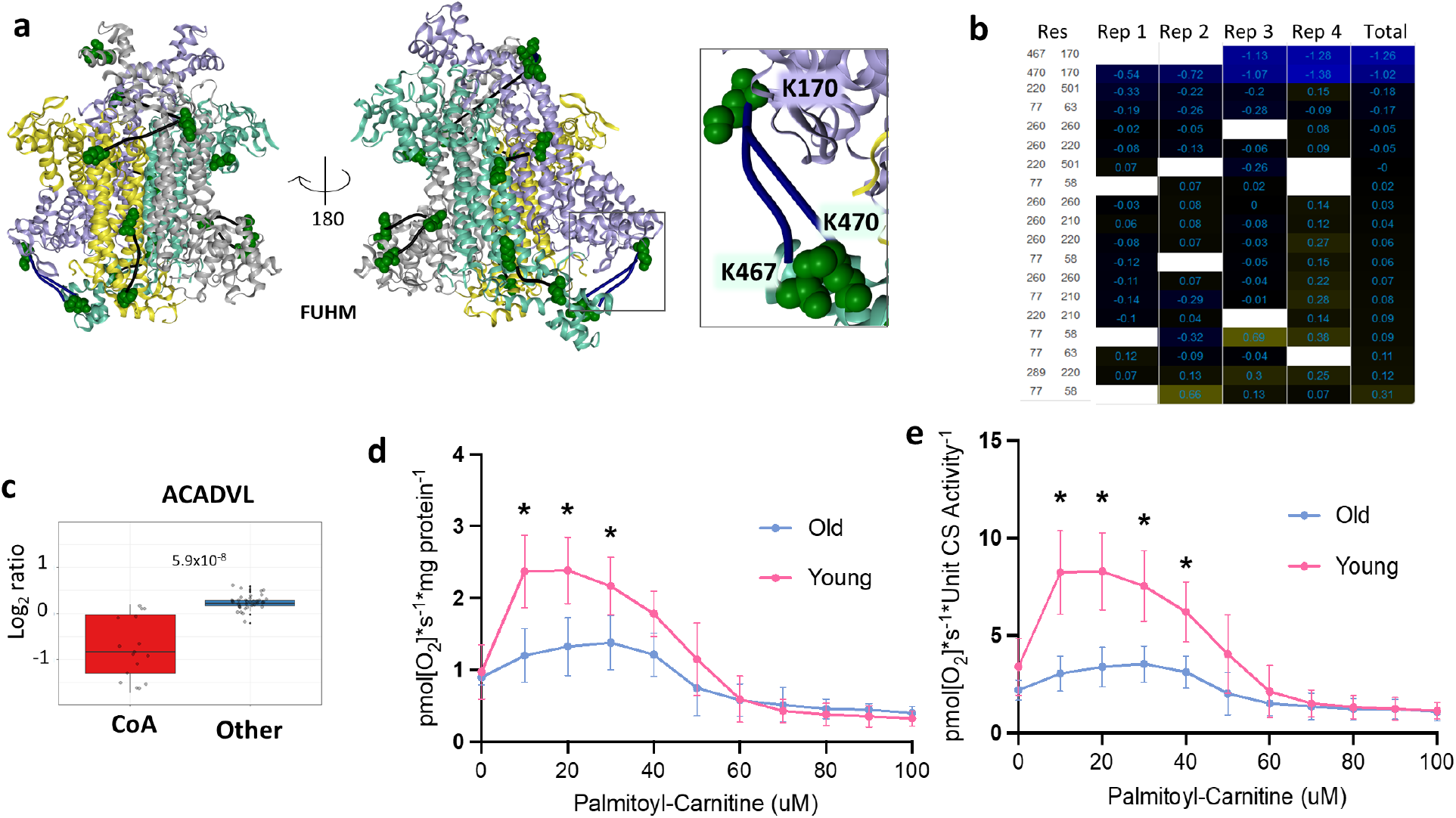
**a**. Fumarate hydratase cross-links mapped to a E.Coli structure. Decreased cross-link levels are shown in the zoomed in square. **b**. Heatmap of log2 ratios of fumarate hydratase cross-links. **c**. Boxplots for Acadvl cross-links based on all 4 biological replicates. **d**,**e**. Palmitoyl-carnitine sensitivity in female F1 mice normalized by mitochondrial protein amount or CS activity.

